# Rehydration rates and the prevalence of xylem-hydration of flowers

**DOI:** 10.1101/255042

**Authors:** Adam B. Roddy, Craig R. Brodersen

**Affiliations:** School of Forestry & Environmental Studies, Yale University, New Haven, CT, USA

**Keywords:** flower, angiosperm, xylem, phloem, water relations, hydraulics

## Abstract

Angiosperm flowers are remarkably diverse anatomically and morphologically, yet they all must satisfy the physiological constraints of supplying sufficient amounts of water and carbon effectively promote pollination. Flowers often occur in the hottest, driest parts of the plant canopy and can face harsh abiotic conditions. Prior evidence suggests that extant species vary dramatically in how water is delivered to flowers, with some evidence that water may be imported into flowers by the phloem. Here we measured midday water potential gradients between flowers, leaves, and stems often phylogenetically diverse species. We further tested the likelihood of xylem-hydration by measuring rates of rehydration after experimentally induced desiccation. There was no significant difference in rehydration rates between leaves and flowers. These results are consistent with xylem-hydration of flowers and suggest that there has been little modification to the mechanisms of water transport despite the diversity of floral form.

## Introduction

Among the angiosperms reproduction has involved the evolution of complex floral structures to attract pollinators, increase outcrossing rates, and protect developing seeds. This critical phase in the life history of a plant can be costly in terms of carbon and water, but these costs can vary widely [1]. Given that most flowers do not assimilate substantial amounts of carbon [2] but may still transpire large quantities of water [3–5], floral transpiration can negatively impact whole plant water balance. Indeed, water lost to floral transpiration can reduce leaf water potential beyond the threshold that induces stomatal closure, thereby suppressing carbon gain and further compounding the costs of reproduction [6–8].

Because of the negative effects of floral water loss and the high carbon invesment into building and maintaining flowers, these costs of reproduction may have driven selection for physiologically cheaper flowers. At a broad phylogenetic scale, floral hydraulic traits vary substantially among lineages [9]. Compared to ANITA grade and magnoliid flowers, monocot and eudicot flowers have lower whole-flower hydraulic conductance, minimum epidermal conductance, and fewer stomata [5]. This trend is in contrast to leaves, which have evolved traits facilitating higher rates of transpiration [10]. This disparity between leaf and flower hydraulic architecture suggests that limiting water loss from floral structures may have been critical in the evolution of their large, morphologically complex flowers.

Furthermore, some evidence suggests that the pathways of water entry into flowers vary substantially among species. Flowers of some ANITA grade and magnoliid species exhibit water potentials consistent with water delivery by the xylem (i.e. flower water potentials more negative than stems and leaves) [11,12], but flowers of some eudicots maintain higher (i.e. less negative) water potentials than leaves. These trends have been used to suggest that they may be hydraulically isolated from the stem xylem and hydrated instead by the phloem [13,14]. The difference in water potential between flowers and vegetative structures can be quite dramatic; petals of cotton plants experiencing drought can maintain water potentials 3 MPa higher than subtending bracts connected to the stem less than one centimeter from the petals [13]. How such large water potential gradients are maintained is unclear, yet may be linked to variation in the pathways of water entry into flowers.

The possiblity of two fundamentally different mechanisms of delivering water to flowers-hydration by the xylem versus the phloem–is appealing because of the potential physiological differences between these two strategies and because of their implications for floral evolution. Long extinct, early angiosperm flowers are thought to have evolved as highly modified leaves, consistent with xylem-hydration of basal angiosperm flowers [11,12]. A transition to phloem-hydration could be beneficial if it helps to buffer flower water potential from variation in plant water potential. Phloem-hydration could result from a combination of reduced transpiration rates and xylem dysfunction. Whether the phloem could supply enough water to maintain turgid, showy flowers given the high hydraulic resistance of the phloem is unclear. Many flowers have lower stomatal densities than leaves [5,15,16], which might allow floral transpiration rates during anthesis to be low enough that water supplied by the phloem and water stored in floral hydraulic capacitors would be sufficient to meet the demands of transpiration. However, while flowers have much higher hydraulic capacitance than leaves [14], they also have significantly higher minimum epidermal conductances [17]. Xylem disconnection between the stem and the flower-due either to discontinuity in the receptacle [18] or to occlusion of the xylem [19]-could physiologically isolate petals from other floral organs and from the stem xylem, allowing petal water potential to vary widely and independently of leaf and stem water potentials.

Data supporting this hypothesis, however, are lacking. To date, water potentials have been measured on flowers of only nine species. Chapotin et al. [14] report water potentials of flowers and leaves of three tropical trees, but for one species flowers and leaves were measured on different individual plants, and no measurements of stem water potentials were made. Inferring directions of water flow from flower and leaf water potentials without measurements of stems is problematic because flowers may have water potentials intermediate between stems and leaves, consistent with xylem-hydration. Indeed, this has been shown in *Illicium* and *Magnolia* flowers, which suggests that these flowers remain hydraulically connected to the stem xylem [11,12]. Although flowers of *Magnolia grandiflora* generally have lower water potentials than stems, inner whorl tepals maintain higher water potentials than stems, which is the only example of floral structures maintaining higher, less negative water potentials than stems [12]. While Trolinder et al. [13] showed that petals can remain significantly more well hydrated than both bracts and leaves, water potentials of stems were not reported, making interpretation of their results difficult.

Thus, the lack of water potentials measured simultaneously on stems, leaves, and flowers hinders our understanding of the potential variation in pathways for water entry into flowers and of floral hydraulic architecture more generally. Here, we report midday water potentials of flowers, leaves, and stems from ten species spanning most of the extant phylogenetic diversity of the angiosperms. We also combine measurements under natural conditions with measurements on slowly desiccating, excised shoots to estimate both the natural variation of midday flower water potential and the magnitude of water potential gradients between flowers and stems under extreme drought conditions. Additionally, these excised shoots were allowed to rehydrate and their water potentials remeasured after 3-4 hours to determine whether and at what rates flower water potentials can recover from declines in water content.

## Methods

Plants growing in the Marsh Botanical Garden (New Haven, CT, USA) and the Arnold Arboretum of Harvard University (Roslindale, MA, USA) were sampled in the spring and summer 2017. These included two *Rhododendron* hybrids, one a likely cross between *Rhododendron catawbiense* and *Rhododendron ponticum* and the other a cultivar in subgenus *Azaleastrum* that has a double corolla (referred to as *Rhododendron catawbiense x ponticum* and *Rhododendron* subg. *Azaleastrum,* respectively), as well as *Magnolia x loebneri,* which is a cross between *Magnolia kobus* and *Magnolia stellata.* Because of differences in floral phenology, species were sampled opportunistically as flowers became available for measurement.

In all experiments, samples were sealed into thermocouple psychrometer chambers within five seconds of excision (Merrill Specialty Equipment, Logan, UT, USA). Within ten minutes of sampling, chambers were triple-bagged in the laboratory, and submerged in a water bath maintained at 25°C for five to seven hours, at which time sequential water potential measurements had stabilized. Water potentials of all structures were made using thermocouple psychrometers interfaced to a CR6 datalogger via an AM16/32B multiplexer (Campbell Scientific, Logan, UT, USA). Measurements of the microvolt output from the psychrometers were converted to MPa using sensor-specific calibration curves generated from measurements of eight NaCl solutions of known water potential [20].

Midday water potentials were measured between 1300 and 1500 hrs on each day from at least three individuals of each species, with the exception of *Clematis montana* var. *rubens,* for which only one individual was available. In the drydown and rehydration experiments, flowering shoots were collected in the morning and immediately enclosed in sealed, humidified plastic bags. After 2-3 hours of equlibration in the plastic bags, initial water potentials were measured. Flowers and leaves were sampled by excising two 6-mm diameter discs of each tissue from midway down the length of the leaf, petal, or tepal and from midway between the midrib and margin, avoiding major veins if possible. The newest, fully expanded leaves on the same shoot as the flower were chosen. Short (~1 cm length) stem segments were excised from below the leaves. All samples were enclosed in thermocouple psychrometer chambers immediately after sampling. In the rehydration experiment, the cut surface of each shoot was placed in distilled water and the shoot allowed to rehydrate for 3-4 hours, at which time water potentials of each structure were resampled. In species with unfused corollas, adjacent tepals or petals of the same flower were sampled, and for species with fused petals, separate but adjacent flowers were sampled. Stem samples after rehydration were taken from just below the sampled leaves, avoiding the approximately 1-cm segment that had been sitting directly in water. This sampling scheme for leaves and flowers assumed that adjacent flowers (or leaves) had the same water potential.

We calculated tissue-specific rehydration rates as:

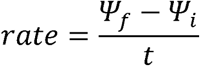

where Ψ_i_ and Ψ_f_ are the water potentials immediately prior to and following rehydration, respectively, and *t* is the time (hours) the sample was allowed to rehydrate. The absolute rate of water potential recovery depends on the water potential gradient between source (approximately 0 MPa for pure water) and the tissue water potential, Ψ_i_. The slope of the relationship between the rehydration rate and Ψ_i_ is the intrinsic time constant of rehydration (τ; hr^−1^). This time constant was calculated for leaves and flowers of each species and compared using a paired t-test.

Because we are not interested in statistical comparisons of water potentials of the same structures between species but rather in the water potential differences between structures within each species, we performed separate mixed-effect ANOVA modeling for each species. For each model, time of day and structure were treated as fixed effects and date and individual as random effects. All analyses were performed in R [21]

## Results

Of the ten species for which there were measurements of midday water potentials, four of them had flower water potentials more negative than stem water potentials, and four of them had flower water potentials indistinguishable from stem water potential (Figure 1). Only two species, *Clematis montana* var. *rubens* and *Weigela coraeensis* had flower water potentials consistently higher (i.e. less negative) than stem water potentials, gradients which have been used previously to argue for phloem-hydration of flowers.

**Figure 1.**
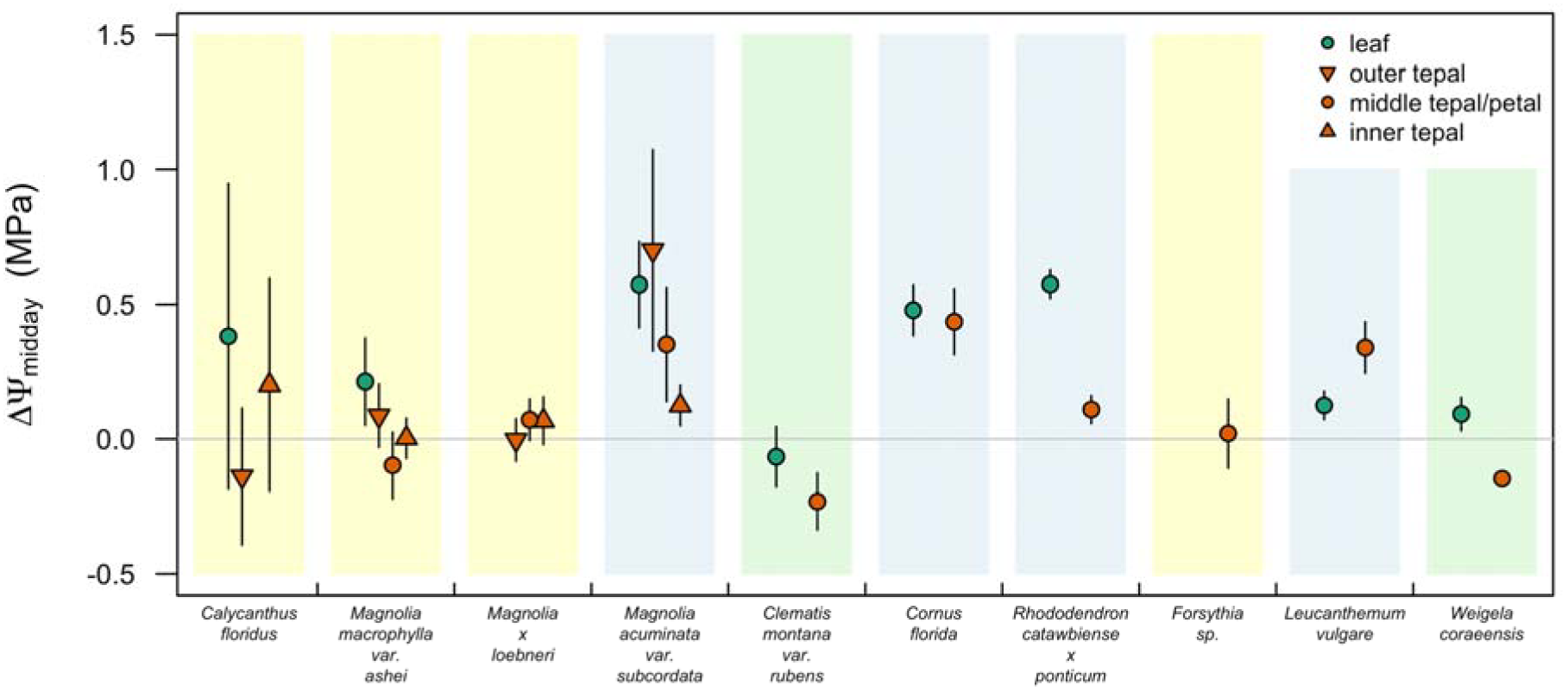
Midday water potential gradients (ΔΨ_stem-leaf_ or ΔΨ_stem-flower_) for ten species measured under natural conditions. Different floral structures are differentiated by different symbols. The grey, horizontal line represents the condition when Ψ of the structure is equivalent to Ψ_stem_ (i.e. ΔΨ = 0 MPa). Positive values indicate that leaf or floral structures have Ψ lower than stems and negative values indicate that leaf or floral structures have Ψ higher than stems. Shading indicates presumed hydration pathway based on water potential gradients (blue: xylem-hydrated; green: phloem-hydrated; yellow: equivocal). Points and error bars represent mean ± s.e.

Of the four species that had flower water potentials close to stem water potential, two of these were precociously flowering species (*Magnolia* x *loebneri* and *Forsythia* sp.) that flower early in the spring when vapor pressure deficits are low and before leaves have flushed. These species may, therefore, not compete with leaves for water. One of these species, *Calycanthus floridus,* has been shown previously to have whole-flower water potentials more negative than stems (Roddy et al., in press), suggesting that while the overall Ψ_stem-flower_ gradient may drive water flow towards flowers, there may be intrafloral variation in water potential gradients between individual tepal whorls.

To determine the ranges of water potentials and the rates of rehydration, we allowed excised flowering shoots to dry on the bench and sampled water potentials periodically over time (Figure 2a). The driest flowers measured of each species showed signs of necrosis, having shriveled and begun turning brown. Yet, mean water potentials of flowers from this experiment never exceeded −1.5 MPa (Figure 2a), and the species with the lowest mean Ψ_*i,flower*_ were tepals of *Calycanthus floridus,* bracts of *Cornus florida,* and petals of *Syringa pubescense.* Mean Ψ_*i,flower*_ of other species were all above −1.0 MPa. Ψ_*i,leaf*_ was generally lower than Ψ_*i,flower*_ for most species.

**Figure 2.**
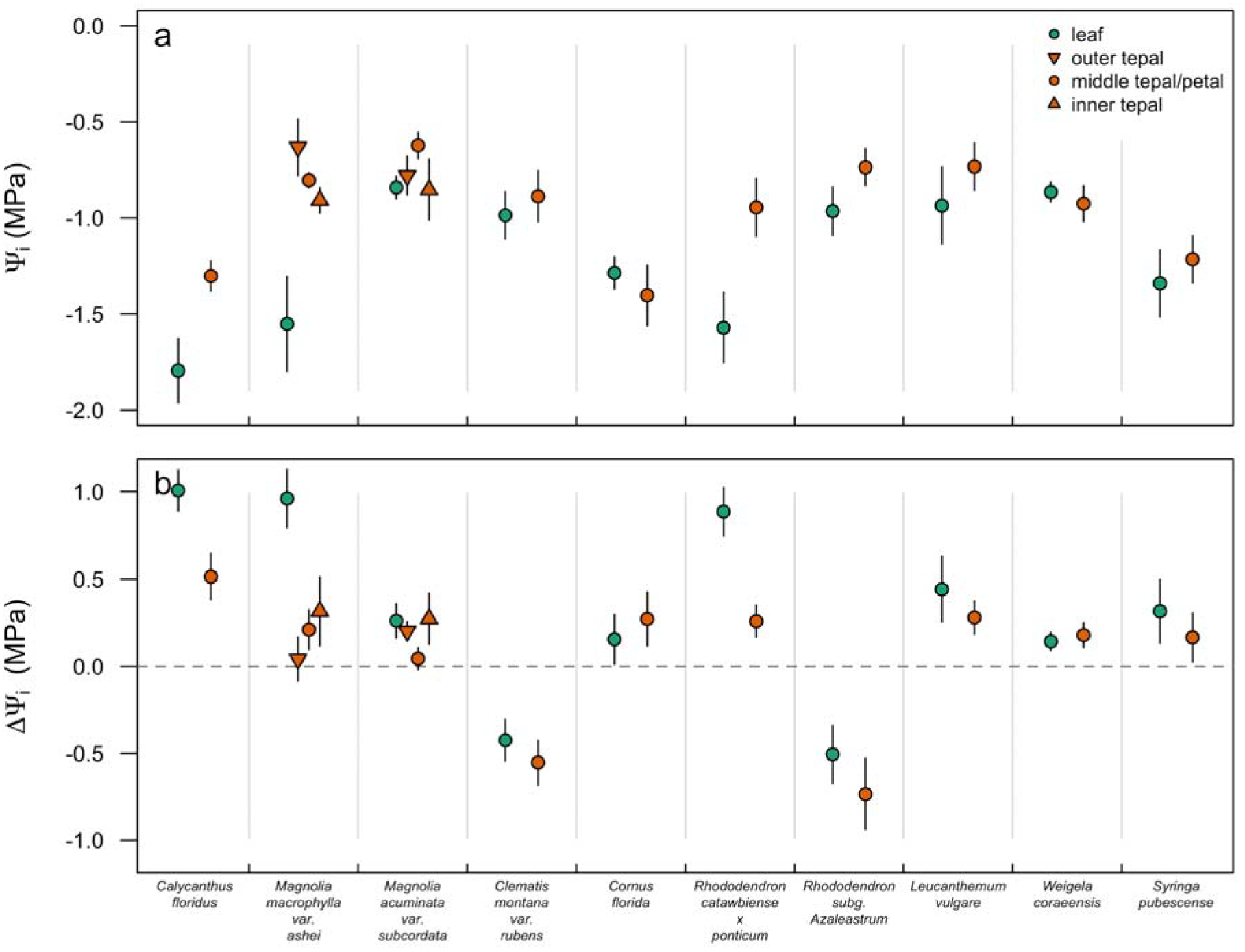
(a) Leaf and flower water potentials and (b) stem-leaf and stem-flower water potential gradients after bench drying and prior to rehydration. The dashed, horizontal line represents the condition when Ψ of the structure is equivalent to Ψ_stem_ (i.e. ΔΨ = 0 MPa). Points and error bars represent mean ± s.e.

However, mean ΔΨ_i_ never exceeded 1 MPa, indicating that leaf and flower Ψ remained very close to Vstem even during benchtop dehydration (Figure 2b). In only five often species was mean ΔΨ_i_ higher than ΔΨ_*middway*_ (*Calycanthus floridus, Magnolia macrophylla, Rhododendron catawbiense* x *ponticum, Leucanthemum vulgare, Weigela coraeensis).* In two species, *Clematis montana* and *Rhododendron* subg. *Azaleastrum,* leaves and flowers remained more well hydrated than stems during dehydration.

More useful information on the effects of water potential declines on hydraulic functioning comes from the rehydration phase of the drydown experiment (Figure 3). For all structures of all species, Ψ_*i*_ was a strong predictor of rehydration rate; samples allowed to desiccate longer with lower Ψ_*i*_ had faster rates of water potential recovery. There was little variation among species and structures in the relationship between Ψ_*i*_ and rehydration rate. To quantify this relationship, we calculated the slope, τ, which is the intrinsic time constant of rehydration. τ did not differ significantly among leaves and flowers (t = 1.64, df = 9, P = 0.14; Figure 4).

**Figure 3.**
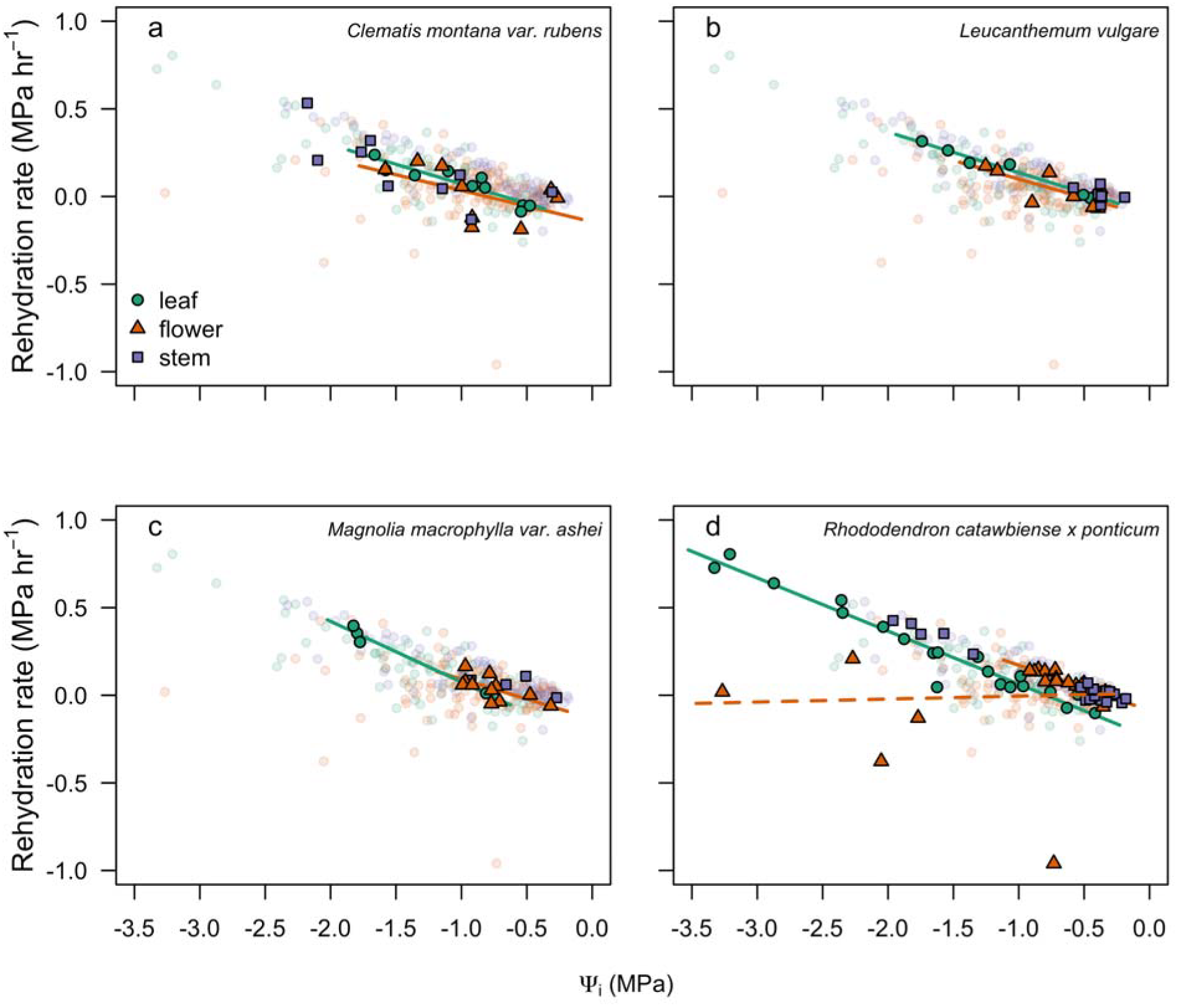
Rehydration rates of leaves, flowers, and stems as a function of initial water potential for (a) *Clematis montana* var. *rubens,* (b) *Leucanthemum vulgare,* (c) *Magnolia macrophylla* var. *ashei,* and (d) *Rhododendron catawbiense xponticum.,* with data for all species in lighter colors. Species-specific regression lines for leaves and flowers are shown. In (d), the solid line for flowers represents only points with initial water potentials greater than −1 MPa, while the dashed line represents all flowers of this species.

**Figure 4.**
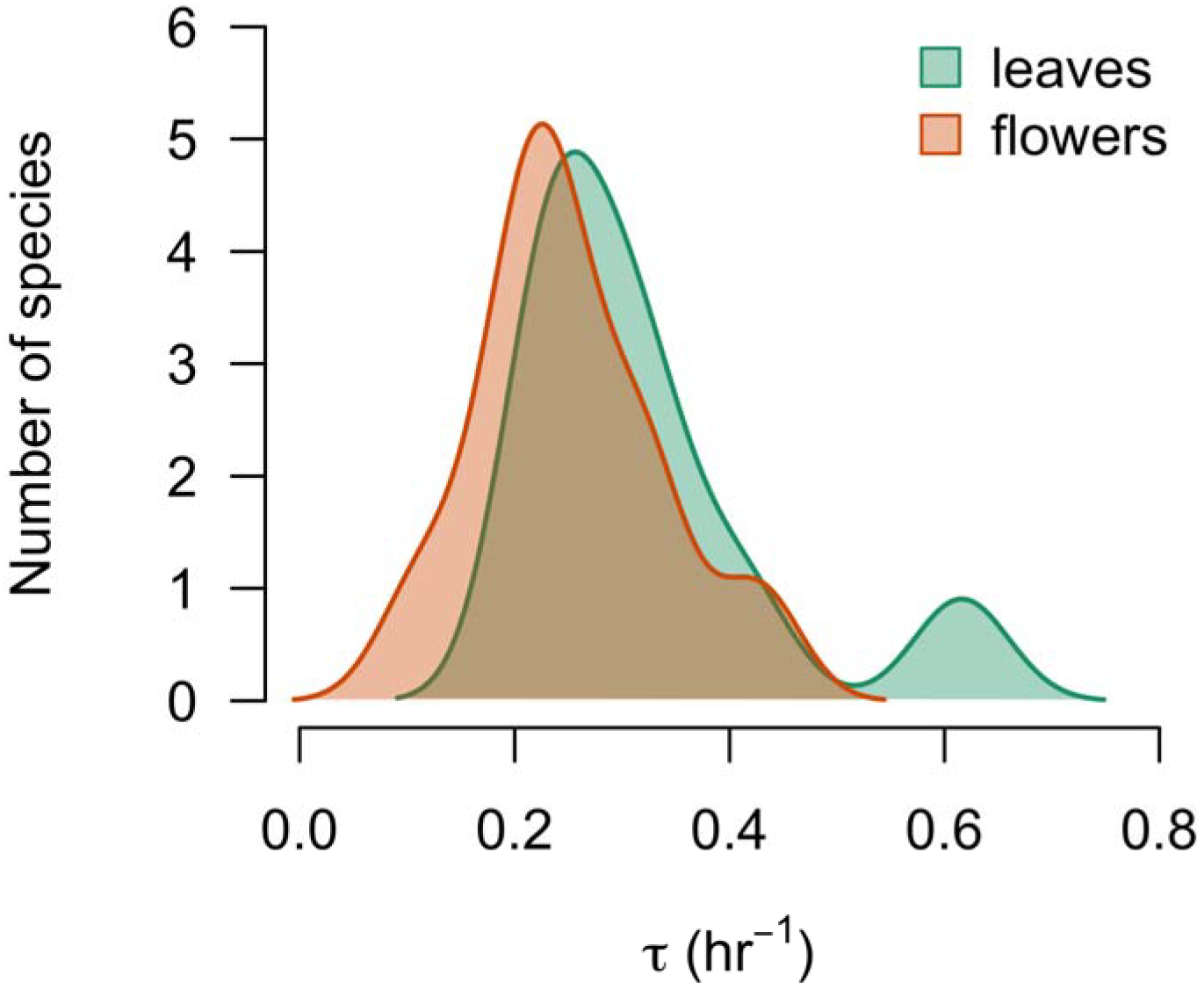
The time constant of rehydration (τ, the slope of rehydration rate versus water potential as shown in Figure 3) did not differ between leaves and flowers.

## Discussion

In contrast to previous reports, water potential gradients between flowers and stems suggest that flowers of many species remain hydraulically connected to the stem xylem during anthesis. Results from the rehydration experiment further corroborate this result. Together these experiments help to clarify the dynamics of water potential gradients in flowering shoots under natural conditions and during experimental desiccation and rehydration cycles.

The direction of water potential gradients between stems and flowers has been surprisingly unclear, with some reports suggesting that flowers may not be hydraulically connected to the stem xylem during anthesis. Reports of flowers having higher water potentials than leaves have been used to suggest that flowers may be hydrated by the phloem [13,14], while other reports have shown that water potentials of flowers are more negative than stems, suggesting that flowers remain hydrated by the stem xylem during anthesis [11,12]. While a single transition from xylem- to phloem-hydration is not necessarily expected, the apparently strong phylogenetic signal in the pathways of water entry to flowers is reinforced by similarly strong phylogenetic signal in other floral hydraulic traits [5]. Our data strongly suggest that most flowers–even those of eudicots, which are purported to be phloem-hydrated-may remain hydraulically connected to the stem xylem. Indeed, ΔΨ_stem-flower_ is often in the same direction as ΔΨ_stem-leaf_ though the magnitude of ΔΨ_stem-flower_ is lower (Figure 1). Therefore, previous data used to show phloem-hydration of flowers are consistent with our results for xylem-hydrated flowers.

However, among some species, it is certainly possible that ΔΨ_stem-flower_ may be negative, which would allow water to flow from the flower to the stem. This occurred among two magnoliids *(Calycanthus floridus, Magnolia macrophylla* var. *ashei),* the basal eudicot *(Clematis montand),* and one of the eudicots *(Weigela coraeensis).* Even with these reverse water potential gradients, how much water may flow from flowers to stems depends upon the resistance in the hydraulic pathway. In this case, flowers may actually supply water to the stem, as do some fruits [22]. Athough the relative contributions of the various resistances in the hydraulic pathway into and through flowers have not yet been quantified, measurements of whole-flower hydraulic conductance suggest that the hydraulic resistances can be high, but not substantially higher than in leaves [5].

The midday water potential gradients reported here also suggest that flower hydraulic architecture may differ between species that flower before or after leaf out. Two of the species measured here were precociously flowering, producing flowers prior to leaf flush *(Magnolia* x *loebneri* and *Forsythia* sp.). Flowers of both of these species had water potentials equal to Ψ_stem_ (Figure 1). Without the need to compete for water with co-occurring leaves, Ψ_flower_ in these species may not need to decline much below Ψ_stem_ in order to drive water flow into the flower. Although precocious flowering has been hypothesized as a way to eliminate competition for water between flowers and leaves, even leaves on the same branch may not compete with each other for water [23], suggesting that there may be little or no competition for water between leaves and flowers. The hydraulic architecture of precocious flowers may differ in other ways from flowers that co-occur with leaves. For example, precocious flowers appear earlier in spring, when atmospheric conditions are cooler and more humid, which limits their transpiration rates [24]. Furthermore, in ring-porous species current-year vessels in the stem bole are mature only once leaves are mature [25], suggesting that the water used by precocious flowers may be provided by localized stem water storage.

The water potential gradients reported also aid in interpreting the role of embolism formation and spread in flowers. Zhang and Brodribb [26] recently reported that water potentials at 50% loss of xylem function in flowers of four species ranged from −2 to −4 MPa, while leaves of the same species ranged from about −1.5 to −7 MPa. The extent to which embolism may influence flower function, phenology, and floral longevity is unclear. Recent evidence suggests that *Calycanthus floridus* flowers rarely encounter water potentials low enough to induce embolism under natural conditions (Roddy et al., in press). Here, we show the lowest midday Ψ_flower_ measured was −1.63 MPa in an outer tepal of *Magnolia acuminata* var. *subcordata.* Thus, it is unlikely that the flowers measured in the present study experienced embolism at midday. Even when shoots had been excised and allowed to desiccate, Ψ_flower_ of almost all species remained higher than −2 MPa. Only petals of *Rhododendron catawbiense* x *ponticum* displayed water potentials substantially below −2 MPa, and these petals did not rehydrate, suggesting that there may have been embolism-induced hydraulic failure or other structural damage to outside-xylem pathways that prevented rehydration. While water loss and the threat of desiccation impact floral display [6]8], our results suggest that under natural conditions, flowers rarely encounter embolism.

Flowers and leaves differed little in their rates of rehydration. While leaf water potentials tended to decline more than flowers during the desiccation experiment, leaves and flowers followed similar rehydration trajectories with no difference in their intrinsic rates of rehydration (Figures 3, 4). With the exception of one *Rhododendron* species, flowers rehydrated just as quickly as leaves for a given initial water potential, suggesting that their lower vein densities and hydraulic conductances did not hinder their capacity to recover from desiccation-induced water potential declines. In contrast to the other species, rehydration rates of *Rhododendron catawbiense* x *ponticum* petals did not follow the same trajectory as leaves or flowers of other species when the initial water potential was below approximately −1 MPa (Figure 3d). Between 0 and −1 MPa, however, this species showed rehydration patterns consistent with the other species studied, suggesting that they may have suffered failure in the hydraulic pathway. Importantly, though, while the rate of water potential recovery did not differ among flowers and leaves (Figure 4), because flower water potentials did not decline as much as leaves, the absolute change in water potential was less in flowers.

Under natural conditions, water potentials of flowers during anthesis deviate little from stem water potentials, with ΔΨ_stem-flower_ rarely exceeding 0.5 MPa, and only in some species were reverse water potential gradients observed (Figure 1). While these Ψ gradients cannot unequivocally determine whether flowers are hydrated by the xylem or by the phloem, the prevalance of positive ΔΨ_stem-flower_ among species is consistent with xylem-hydration of flowers, even among the eudicots. Given that flowers lose turgor at higher water potentials than leaves [12], minimizing ΔΨ_stem-flower_ may be critical to preventing turgor loss. Furthermore, these results suggest that flowers can rehydrate as rapidly as leaves. Unlike leaves, however, which must remain turgid to continue assimilating carbon, it is possible that wilted flowers may still attract pollinators as long as ovary water potentials remain high. Although the pathways for water movement into flowers remain unclear, our measurements of midday water potentials and of rehydration dynamics do not rule out the possibility of xylem hydration. Indeed, xylem hydration of flowers is certainly possible, and the apparent dichotomy between xylem-hydration of basal angiosperm flowers and phloem-hydration of eudicot flowers may very well be spurious.

## Acknowledgments

K. Richardson, M. Dosmann, and F. Rosin provided access and support at the Arnold Arboretum. Funding was provided by the Arnold Arboretum of Harvard University and the Yale Institute for Biospheric Studies.

